# BK Channels Orchestrate Cardiac Homeostasis Through Mitochondrial Uncoupling Proteins

**DOI:** 10.64898/2026.05.20.726286

**Authors:** Shubha Gururaja Rao, Nishi Patel, Neel Patel, Kajol Shah, Ahmed Hussain, Satish K. Raut, Swaiman Singh, Devasena Ponnalagu, Sankar Adya, Federica Accornero, Andrew Kohut, Harpreet Singh

## Abstract

BK channels, coded by the *Kcnma1* gene, integrate voltage and intracellular Ca^2+^ signals and are recognized for their roles in smooth muscle and neuronal excitability. However, their contribution to baseline cardiac physiology remains poorly defined. Here we uncover a fundamental function for BK channels in maintaining normal cardiac performance, independent of pathological stress. Using non-invasive echocardiography, transcriptional profiling, and mechanistic analyses, we demonstrate that *Kcnma1* deletion disrupts ventricular function, and remodels metabolic and stress-response pathways. Transcriptomic profiling revealed selective downregulation of mitochondrial uncoupling proteins (UCPs) and suppression of the PGC-1α/FOXO3a axis, without broad loss of oxidative phosphorylation components. Enhancing UCP expression restored cardiac performance, indicating that mitochondrial uncoupling and redox control constitute key downstream effectors of BK signaling. Together, these results identify a physiological role for BK channels in maintaining myocardial function and define a mitochondrial BK-UCP axis, critical for cardiac homeostasis.

## Introduction

Large-conductance, voltage- and calcium (Ca^2+^) -activated potassium (BK, Slo, MaxiK) channels are unique among K^+^ channels because they uniquely integrate membrane depolarization with local intracellular Ca^2+^ microdomains to regulate excitability across a wide range of tissues(*1-3*). A single pore forming α-subunit is encoded by a gene, *Kcnma1(4)*, gives rise to an unexpectedly diverse array of channel behaviors. Despite originating from one gene, BK channels exhibit remarkable heterogeneity in gating kinetics, pharmacological sensitivity, subcellular distribution, and physiological roles(*5, 6*). This diversity arises from extensive post-transcriptional regulation (including alternative pre-mRNA splicing), post-translational modifications, and co-assembly with auxiliary β- (*Kcnmb*1-4) and γ- (LRRC26/38/52/55) subunits, which tune BK channel voltage and Ca^2+^ sensitivity, trafficking, and tissue-specific functions(*4, 7*). Although these molecular mechanisms generating BK complexity are increasingly well characterized, their functional significance across organ systems remains incompletely understood.

BK channels display several biophysical and physiological features that position them as high-gain regulators of cellular activity(*4*). Their unusually large single-channel conductance which is in order of 10-20 times greater than most other K^+^ channels(*3, 4*) enables them to rapidly repolarize membranes and counteract excitatory stimuli. In smooth muscle, a single Ca^2+^ spark can recruit ∼15 BK channels, producing ∼20 mV of hyperpolarization that closes voltage-dependent Ca^2+^ channels and thereby relaxes the muscle(*8*). In the nervous system, BK channels are widely expressed in cortical and hippocampal neurons, where they shape action-potential duration, firing frequency, spike timing, and neurotransmitter release(*9, 10*). In visceral organs, such as the urinary bladder, BK channels govern detrusor contractility and promote controlled filling and voiding cycles(*11*). Consistent with their multifunctional character, BK channels have been detected not only in the plasma membrane but also in intracellular compartments including the nucleus, endoplasmic reticulum, secretory vesicles, and mitochondria, where they modulate organelle-specific Ca^2+^ and redox dynamics mitochondria(*1, 4, 12-15*).

In the cardiovascular system, the role of BK channels is complex and compartment-specific(*1, 12, 14*). Although conventional electrophysiology often fails to detect BK current at the surface membrane of adult ventricular myocytes, converging evidence highlights two prominent cardiac axes for BK function(*1*). First, within the cardiac conduction system, BK channels modulate sinoatrial node (SAN) automaticity and autonomic responsiveness(*16, 17*). Pharmacological blockade slows SAN firing and blunts adrenergic rate acceleration, while genetic models across mammals, and *Drosophila* consistently link *Kcnma1* dysfunction to sinus bradycardia and arrhythmogenicity(*16-20*). Second, within the working myocardium, a mitochondrial pool of BK channels (mitoBK) has been implicated in cardioprotection, particularly during ischemia– reperfusion (IR) stress(*7, 18, 21-25*). Studies using genetic models and BK-channel openers demonstrate that mitoBK activity can limit ROS generation, stabilize mitochondrial Ca^2+^, reduce permeability-transition susceptibility, and ultimately decrease myocardial infarction size(*7, 18, 25-28*).

However, an overwhelming majority of our current understanding of cardiac BK function arises from disease-focused IR models(*7, 18, 19, 25, 26, 28*). While instrumental in establishing the cardioprotective role of mitoBK channels, these approaches offer only a narrow view and leave the physiological, baseline function of BK channels in the intact heart poorly defined. Similarly, much of the pharmacological evidence remains confounded by the lack of channel specificity of early BK modulators. For example, prior work using fast-acting BK blockers demonstrated reversible cardiovascular effects in rodents(*7, 20-27*) underscoring the importance of BK channels in real-time cardiac and vascular regulation but without resolving whether these effects arise from conduction system alterations, vascular tone changes, mitochondrial modulation, or integrated whole-organ interactions.

Given these knowledge gaps, it remains unknown whether BK channels are required for normal cardiovascular homeostasis, independently of injury paradigms. To address this, we employed non-invasive echocardiography and complementary mechanistic assays to interrogate the role of BK channels in cardiac function under physiological conditions. By expanding beyond traditional IR-based frameworks, our study provides a broader and more integrated understanding of BK-mediated cardiac adaptation via the regulation of mitochondrial uncoupling proteins.

## Methods

### Animals

Experiments were performed using 2-3-month-old wild type and *Kcnma1*^*-/-*^ C57BL/6 mice. The experimental procedures were designed in accordance with the National Institutes of Health and American Association for the Accreditation of Laboratory Animal Care (AAALAC) guidelines and approved by the Drexel University College of Medicine Institutional Animal Care and Use Committee (IACUC), Philadelphia, PA, and the Ohio State University IACUC, Columbus, OH.

### Cardiac functional analysis

A high-frequency, high-resolution digital imaging platform with linear array technology and Color Doppler Mode for *in vivo* high-resolution micro-imaging was used for echocardiography of mice hearts as per previously published guidelines^35^ (Vevo® 2100 Imaging System, FUJIFILM VisualSonics Inc., Toronto, Canada). Mice were maintained at 37°C on a prewarmed imaging platform. For assessing the cardiovascular function of mice, a high-frequency transducer probe (VisualSonics MS250 with a frequency range of 18-38 MHz) was used as it provides the appropriate resolution and depth of penetration needed.

Mice were anesthetized using 2.5% (*v/v*) isoflurane mixed with oxygen. After achieving adequate anesthesia in mice, anthropomorphic measurements were taken. For echocardiography, hairs from the ventral chest were removed using commercially available hair removal cream (Nair™, © Church & Dwight Co). After securing each mouse to the imaging platform, anesthesia concentration was titrated from 1% to 3% to maintain a minimum heart rate of 400 beats/min. All recordings were acquired and analyzed using Vevo Lab software in consultation with a clinical echocardiographer to ensure optimal image quality.

### Microarray

Adult cardiomyocytes of wild type and *Kcnma1*^*-/-*^ mice were isolated and total RNA was prepared using a commercially available Qiagen RNAeasy kit. RNA was treated with RNase-free DNase I. RNA was quantified on a Nanodrop ND-100 spectrophotometer (NanoDrop Technologies, Wilmington, DE, USA), followed by RNA quality assessment by analysis on an Agilent 2100 bioanalyzer (Agilent, Palo Alto, CA, USA). Fragmented biotin-labeled cDNA (from 100 ng RNA) was prepared using the GeneChip WT Plus kit.

Each Affymetrix gene chip *mouse* array (Affymetrix, Santa Clara, CA, USA) was hybridized with the fragmented and biotin-labeled target (4.5 μg) in 200 μL of hybridization cocktail. Target denaturation was performed at 99 °C for 2 min and then 45 °C for 5 min, followed by hybridization for 18 h. Then the arrays were washed and stained using GeneChip Fluidic Station 450, and hybridization signals were amplified using antibody amplification with goat IgG (Sigma-Aldrich, St. Louis, MO, USA) and anti-streptavidin biotinylated antibody (Vector Laboratories, Burlingame, CA, USA). The chips were scanned on an Affymetrix Gene Chip Scanner 3000, using Command Console Software.

Background correction and normalization were done using Robust Multichip Average with Genespring V 14.9 software (Agilent). A 1.5-fold differentially expressed gene (*p* ≤ 0.05 values) list was generated. The listing of differentially expressed genes and their fold change were loaded into Ingenuity Pathway Analysis (IPA) 5.0 software (Qiagen Inc.) to perform biological network and functional analyses. IPA converts gene sets (with or without expression information) into related molecular networks based on the IPA knowledge database. Core analysis was performed and the genes were categorized based on molecular function, mapped to genetic networks, and ranked by score^36^. The score reflects the probability that a collection of genes equal to or greater than a number in the network could not be achieved by chance alone. A score of more than 10 was used as a cutoff for identifying specific gene networks (n = 3 independent experiments with RNA isolated from mouse hearts).

### Western blot analysis

Ventricular tissue from wild type and *Kcnma1*^*-/-*^ mice were lysed in ice cold RIPA buffer [Tris HCl 50 mM, NaCl 150 mM, EDTA-Na2 1 mM, EGTA-Na4 1 mM, Na3VO4 1 mM, NaF 1 mM, 1% (vol/vol) Nonidet P-40, 0.5% (wt/vol) Na-deoxycholate and 0.1% (wt/vol) SDS, pH 7.4] containing protease inhibitors (1 tablet/50 mL; Roche), and incubated 1 h at 4°C with shaking. Samples were centrifuged for 10 min at 13,000 g and the supernatants were collected. Proteins (100 μg/lane) were separated on 4-20% (*v/v*) SDS/PAGE and transferred to nitrocellulose membranes. Loading was corroborated with Ponceau S staining. Nitrocellulose membranes were blocked with Li-Cor blocking solution at room temperature for 60 min and incubated overnight with antibodies (see in the next paragraph). Membranes were washed thrice with 1x TBS and incubated with 0.01 μg/mL secondary Abs for 60 min at room temperature. After washing three times for 15 mins, membranes were visualized using Odyssey Imaging System.

### Antibodies

UCP1 (Novus biologicals NBP2-20796, used at 1:100), UCP2 Antibody ((G-6): sc-390189, used at 1:100), and UCP3 Antibody (Novus biologicals NBP2-24608, used at 1:100) FoxO3a (Cell signaling D19A7,12829, used at 1:200), Phospho-FoxO3a (Ser253) Antibody (Cell signaling, 9466, used at 1:200) (PGC1α NBP1-04676, 1:100)

### Statistical analysis

For physiologic parameters of interest, four measurements for each animal were taken and averaged. All values were entered into Microsoft Excel, and Student’s t-test was used to compare means for statistical significance. All values are reported as mean ± standard error.

For cell biology, data were analyzed using the Sigma plot. Student’s t-tests or ANOVA were used for analyzing all the data and reported as mean ± standard error of the mean in text. p-values less than 0.05 were considered significant.

## Results

### BK regulates cardiac function

There has been a substantial focus on the role of cardiac BK in cardioprotection, without addressing its physiological role in the heart(*14, 29*). Here we established the role of BK in cardiac function using comprehensive cardiac evaluation using echocardiography(*30, 31*). First, we analyzed available databases for mRNA changes in dilated cardiomyopathy of end stage heart failure with reduced ejection fraction (HFrEF)(*32, 33*). RNA-seq from 51 human left ventricles samples: 37 dilated cardiomyopathy (DCM) and 14 non-failing (NF) showed that expression of BK is significantly (p= 0.01) elevated in male left ventricle (**Supplementary fig. 1**). There was no difference in expression of mRNA between males and females’ left ventricle or end stage heart failure in female DCM patients (**Supplementary fig. 1**). These results indicate that failing male hearts show reprograming of expression of BK channels.

A comprehensive cardiac evaluation of *Kcnma1*^*-/-*^ has indicated intriguing possibilities for the role of BK in cardiac function (**fig. 1)**. LV sections from wt and *Kcnma1*^*-/-*^ mice labeled with Masson trichrome staining revealed a higher degree of fibrosis (**fig. 1A-C**) and an increase in the size of cardiomyocytes (**fig. 1B-C)** in *Kcnma1*^*-/-*^ mice, *implying that absence of mitoBK results in cardiac hypertrophy and fibrosis*. Next, we performed echocardiography on wild-type (wt) and *Kcnma1*^*-/-*^ mice. Echocardiography further indicated dilated ventricles (**fig. 1D**), an increase in LVID (**fig. 1E**), IVSD (**fig. 1F**), and posterior wall (PW) (**fig. 1G**) thickness.

**Figure. 1.**
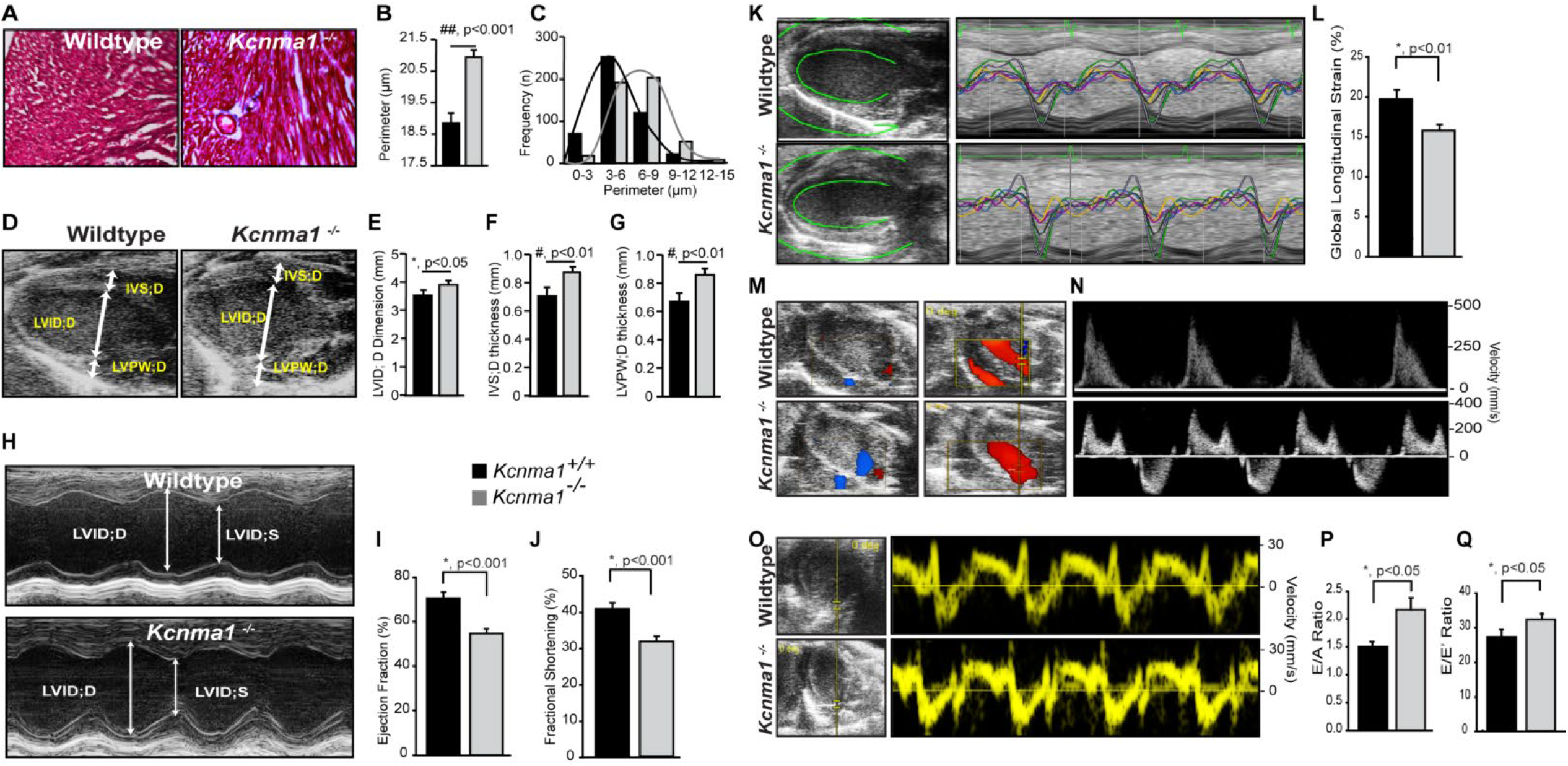
The absence of BK causes cardiac hypertrophy and dysfunction. **A**. Masson trichrome staining. There is a significantly increased collagen deposition (blue) in KO mice heart. **B**. and **C**. Total perimeter and frequency of cardiomyocytes. **D**. Echocardiography showing PSLAX view of wt and KO hearts. **E. F. G**. Wall thickness measured from **D**. and **H. H**. M-mode Echocardiography traces in adult wt and KO mice, respectively. **I**. Comparison of LVEF of wt and KO mice, n=5, p< 0.01. F **J**. Comparison of FS of wt and KO mice, n=5, *, p<0.01. **K**. Strain analysis of WT & KO hearts. **L**. Quantification of global longitudinal strain of wt and KO mice, n=5, *, p<0.01. **M** and **N**. Color Doppler and region of pulse wave Doppler interrogation. Tissue Doppler to derive Tei index. **P**. Comparison of E/A peak early to a late diastolic mitral inflow velocity ratio of WT and KO, n=5, *, p<0.05. **Q**. E/E′ peak early diastolic mitral inflow velocity to tissue Doppler mitral annular velocity ratio of WT and KO, n=5, *, p<0.05.

To evaluate the degree of cardiac dysfunction caused by the increased fibrosis seen in *Kcnma1*^*-/-*^. M-mode measurements of the LV were carried out, as quantitative estimates of LF function are amongst the most valuable prognostic indicators regarding cardiac morbidity and mortality (**fig. 1H-O**). We found that *Kcnma1*^*-/-*^ mice had lower LVEF (51±3 %, n=5) as opposed to 65±5%, n=5 for wt mice (**fig. 1H and I**). LV fractional shortening (LVFS) was significantly (p<0.001) reduced from 39±4%, n=5, for wt mice to 31+2.5%, n=5 for *Kcnma1*^*-/-*^ mice (**fig. 1J**). Global longitudinal strain (GLS) by speckle tracking echocardiography is a key method for evaluation of LV function with enhanced reproducibility compared with LVEF. Strain by speckle tracking echocardiography utilizes two-dimensional grayscale images to evaluate both global and regional functions of the LV(*31*). Cardiac dysfunction was also corroborated by global strain analysis (**fig. 1K and L**), where *Kcnma1*^*-/-*^ mice showed 15±1.5% as compared to 19±2%. Cardiac output, the function of HR, and stroke volume (SV) calculated from echocardiography were significantly reduced from 15±3 to 7±2 mL/min, n=5. We further performed mitral annular plane systolic excursion (MAPSE) as it is known as a surrogate measurement for LVEF in patients(*31*). Our preliminary data show a ∼33% reduction in MAPSE in *Kcnma1*^*-/-*^ mice (**fig. 1M-Q**). We also measured the E(early)/A(late) ratio a marker of LV diastolic filling and found that *Kcnma1*^*-/-*^ has higher E/A ratios, 2.1±0.4 as opposed to 1.17±0.05 for wt mice (**fig. 1P and Q**), and E/E’ ratios, 27±1.8 *vs*. 31±1.5 for *Kcnma1*^*-/-*^ *vs. wt* mice, respectively.

Since BK is vital for maintaining the function of the left heart, we also evaluated the role of BK in the right heart by echocardiography. As shown in **fig. 2**, the right ventricle ejection fraction (RVEF) of *Kcnma1*^*-/-*^ mice was 46±3% *vs*. 63±4% for wt mice (**Supplementary fig. 2A and B**). We also measure pulmonary arterial flow velocities. As shown in **Supplementary fig. 1C-F**, Doppler probing indicated that the peak velocity of *Kcnma1*^*-/-*^ was 360±60 mm/sec as opposed to 595±30 mm/sec for wild-type mice. We also used tricuspid annular plane systolic excursion (TAPSE) analysis which is another two-dimensional measure with which one can assess right ventricular systolic function (**Supplementary fig. 2G and I**). TAPSE was obtained by placing the M-mode cursor through the lateral portion of the tricuspid valve annulus in the apical four-chamber view. *Kcnma1*^*-/-*^ mice showed 0.85±0.1 mm and wild-type mice showed 1.15±0.07 TAPSE (**Supplementary fig. 2G and H**). We also measured the tissue Doppler-derived right ventricular systolic excursion velocity S’ for *Kcnma1*^*-/-*^ and *wt* mice. Similar to TAPSE, tissue doppler systolic velocity of the tricuspid annulus is an additional measure of longitudinal RV systolic performance. Tissue Doppler-derived right ventricular systolic excursion velocity S’ for *Kcnma1*^*-/-*^ *mice was 22*±*2 mm/s and 26*±*1*.*4 mm/s for wt* mice (**Supplementary fig. 2I and J**). Our data from LV and RV indicate global cardiac dysfunction in *Kcnma1*^*-/-*^ mice.

**Fig. 2.**
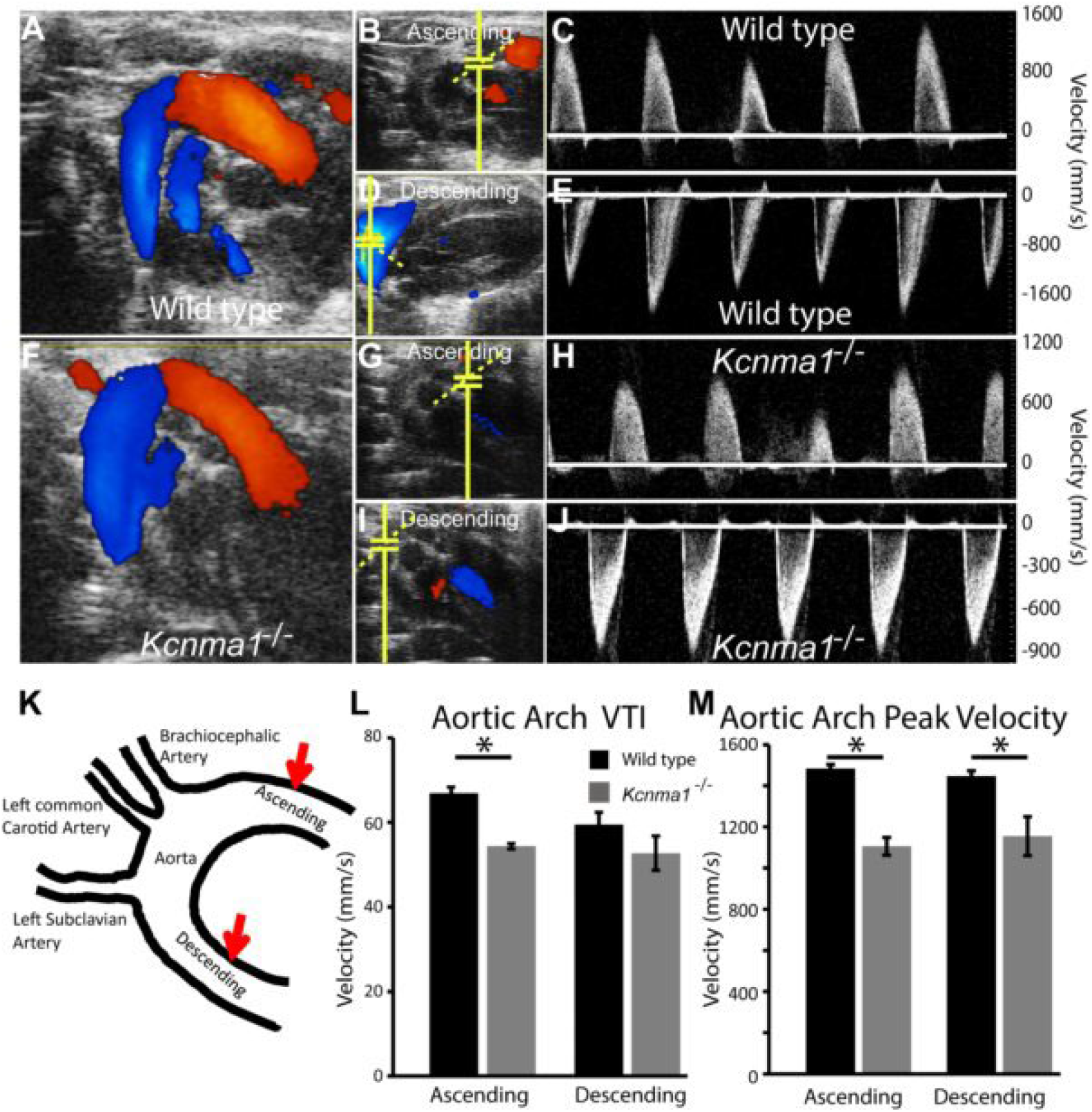
Ablation of BK decreases the aortic flow. **A** and **F**, Color Doppler of ascending and descending aorta, **B, D, G** and **I**, shows region of pulse wave Doppler interrogation; **C** and **E**, pulse wave Doppler flow velocities of the ascending and descending aorta in wt mice. **H** and **J**, pulse wave Doppler flow velocities of the ascending and descending aorta in *Kcnma1*^*-/-*^. **K**. Schematic of aorta, red arrows indicate position of the probe. **L** and **M**. Comparison of VTI and peak velocities of wt and *Kcnma1*^*-/-*^. n=3, *,p< 0.05

### Role of BK in aortic function

Aortic endothelium smooth muscle preparations have been extensively used to investigate the role of K^+^ channels to relax the smooth muscle. The addition of the BK channel opener, NS1619 also results in vasorelaxation. NS-1619 also showed a rightward shift in the concentration-response relationship; however, this compound exhibited similar maximum efficacy at 100 μM under both extracellular K^+^ conditions(*34*). Our experiments implicated the role of BK channels in aortic smooth muscle relaxations by BK channels which corroborates the cellular data(*35-37*). However, along with BK activation, NS-1619 had been shown to also inhibit Ca^2+^ and Cl^−^ channels(*38, 39*). To test the direct role of BK channels in aortic function, we performed Doppler probing of the aorta in wt and *Kcnma1*^*-/-*^ mice. The aortic arch was identified by color Doppler (**fig. 2A** and **F**) and measurements were obtained from ascending and descending arch in wt (**fig. 2B-E**) and *Kcnma1*^*-/-*^ mice (**fig. 2G-J**). Aortic arch VTI and aortic arch peak velocities were measured from **fig. 2C** and **fig. 2E** for wt mice, and **fig. 2H** and **fig. 2J** for *Kcnma1*^*-/-*^ mice. The absence of BK decreased aortic arch VTI for both ascending and descending aorta as compared to mice with BK channels (**fig. 2 K-M**). Maximum peak velocity was also significantly reduced in *Kcnma1*^*-/-*^ mice for both ascending and descending aorta as compared to wt mice. Our results for the first time implicate BK channels directly in the vascular function of the aorta.

### BK channels regulate mitochondrial uncoupling proteins

To elucidate the signaling mechanism involved in BK-mediated cardiac dysfunction, we performed microarray on cardiomyocytes isolated from wt *vs. Kcnma1*^*-/-*^ mice without any pathological or physiological stress. We have discovered that genes involved in cardiac dysfunction showed a significant change in expression (**fig. 3A**). Several genes including TCAP, Cav3, Rrad, Serpine1, Pfkb1, and Fhl2, characterized in HF(*40*) showed a significant change (-2 to 5.5-fold) in expression. We analyzed the profile of all the genes undergoing significant changes and performed in silico analysis for their involvement in cardiac dysfunction. As shown in **fig. 3B**, we discovered that ablation of BK causes augmentation of pathways involved in cardiac hypertrophy (4.19E-06), fibrosis (6.81E-04), arrhythmias, and cardiac necrosis/cell death (7.30E-04).

**Fig. 3.**
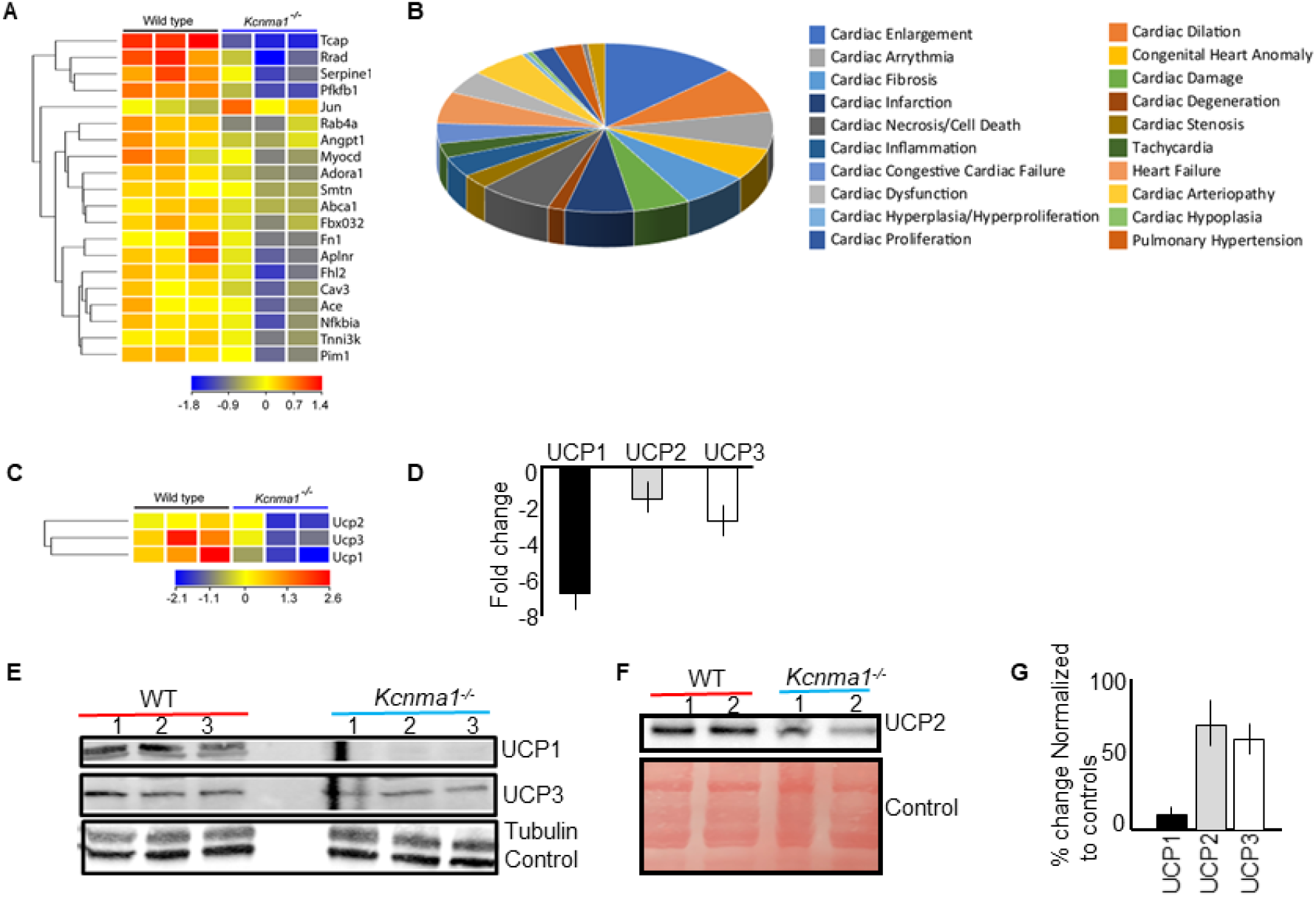
Ablation of BK causes a change in cardiac dysfunction pathways. **A**. Pie chart representing various cardiac dysfunction pathways. Numbers represent a percentage. **B**. Genes involved in cardiac dysfunction showed a change in expression in *Kcnma1*^*-/-*^ cardiomyocytes. **C**. UCP1,2,3 were down-regulated in *Kcnma1*^*-/-*^. **D**. Quantification of UCP1, 2 & 3 mRNA. **E**. Western blot show down regulation of UCP1 (5 fold), and 3 (1.5 fold). **F**. Decrease in levels of UCP2 (2 fold) in *Kcnma1*^*-/-*^. GAPDH and Ponceaus S are controls. **G**. Quantification of F.

Since BK is exclusively localized to mitochondria in adult cardiomyocytes, we also analyzed our array results for mitochondrial genes and found that all three UCPs implicated in cardiac function(*41-43*) were downregulated, and astonishingly UCP1 was 7-fold lower than controls **(fig. 3C** and **D**), We further confirmed the expression of UCP1, 2, and 3 by western blot analysis. Similar to the downregulation of UCP genes, protein expressions of UCP1 and UCP3 were significantly reduced in *Kcnma1*^*-/-*^ mice (**figs. 3E, F, and G**). UCP2 also showed decreased expression but it was not significant. We further evaluated changes in expression of mitochondrial complexes, and we did not observe any difference (**Supplementary fig. 3**). Taking together our microarray and biochemical analysis shows that ablation of BK_Ca_ channels results in the downregulation of UCPs while the core mitochondrial electron transport chain complex levels are unaffected.

### β3-adrenergic receptors agonists reverse cardiac dysfunction in BK knockouts

Since ablation of BK results in cardiac dysfunction by downregulating UCPs, we tested whether increasing expression of UCPs by activating β3-adrenergic receptors using a selective agonist CL-316,243(*44*) will protect the *Kcnma1*^*-/-*^ mice heart. Wild-type mice receiving CL-316,243 compound showed no difference in LVEF (54±2% vs. 59±3%, n=5), but *Kcnma1*^*-/-*^ mice showed significant improvement from 44±3% to 57±4% (n=5) within 4 weeks (**fig. 4A**), reaching the range of wild type LVEF. There was no change in LVEF observed in wild-type or *Kcnma1*^*-/-*^ mice after saline injection (**fig. 4A**). We further probed whether the expression of UCPs changed in *Kcnma1*^*-/-*^ mice by western blots upon CL-316,243 treatment. CL-316,243 significantly enhanced the expression of UCP1 and UCP2 as compared to the saline control (**fig. 4B** and **C**). These results clearly indicate that cardiac dysfunction observed in *Kcnma1*^*-/-*^ mice is mediated by the downregulation of UCPs.

**Fig. 4.**
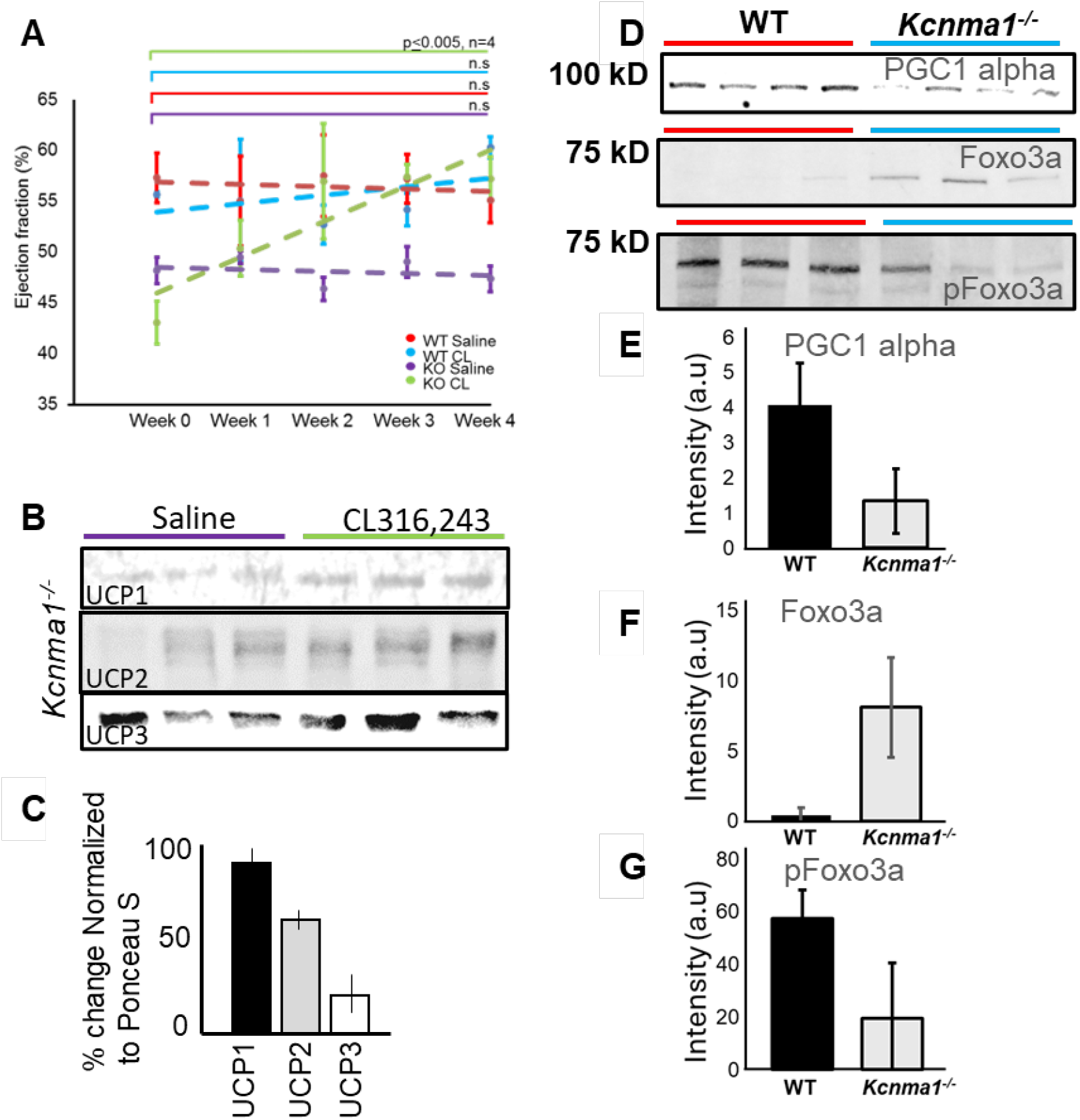
Pharmacological overexpression of UCPs improves cardiac function. **A**. CL316,243 compound (*i*.*p*. 1mg/kg) weekly for four weeks increased the LVEF of BK KO mice (green) but not of mice injected with saline (purple). WT mice injected with CL316,243 (red) or saline (blue) showed no difference in ejection fraction. Our preliminary data indicate that CL316,243 specifically improved the LVEF of KO hearts. There was a small but insignificant improvement in WT (light blue). N=4 in all groups. **B**. Cardiac lysates of *Kcnma1*^*-/-*^ mice from A (with and without CL compound after week 4) were probed for expression of PGC-1α, pFoxo3a and Foxo3a. Expression of PGC-1α and pFoxo3a increases but Foxo3a does not show any significant change in expression. **C**. Quantification of UCP1, 2 and 3 expression from **B** and normalized to Ponceau-S stained membrane. **D**. Western blot shows a significant reduction in the expression of PGC-1α and pFoxo3a in BK knockout animals but increase in the expression of Foxo3a levels. Quantification is provided for PGC-1α in **E**, pFoxo3a in G and Foxo3a in **F.**

In order to elucidate how BK is changing the expression of UCPs, we probed proteins that regulate the expression of UCPs. Peroxisome proliferator-activated receptor-γ coactivator-1α (PGC-1α) is the mitochondrial master transcriptional coactivator that coordinates mitochondrial biogenesis and oxidative metabolism. One of its most physiologically important functions in this context is the induction of UCPs, which modulate mitochondrial efficiency, heat production, and reactive oxygen species (ROS) handling(*41-47*). We found that PGC-1α is 3-fold down-regulated (**fig. 4D and 4E**) in *Kcnma1*^*-/-*^ mice. Expression of proteins in cardiac hypertrophy is also governed by Foxo3a(*48-50*), and we found that though Foxo3a expression is increased, the activated Foxo3a (phosphorylated) is down-regulated in *Kcnma1*^*-/-*^ (**fig. 4D and F,J**). Our results are the first evidence that the expression of BK modulates the expression of transcription factors, pFoxo3a and PGC-1α.

## Discussion

BK channels are uniquely poised to couple intracellular Ca^2+^ signals to membrane hyperpolarization owing to their exceptionally high unitary conductance and dual gating by voltage and Ca^2+^(*51*). They thereby serve as rapid, negative-feedback regulators of excitability across excitable and non-excitable tissues, including neurons, vascular smooth muscle, and cardiac pacemaker and fibroblast populations(*52*). In the cardiovascular system, BK channels in vascular smooth muscle promote vasodilation and set myogenic tone, while in the heart they have been implicated in rate control through sinoatrial node (SAN) mechanisms and in cytoprotection through a mitochondria-localized pool in adult ventricular cardiomyocytes(*16, 17, 19, 20*). BK channels (and their mitochondrial counterparts, “mitoBK”) have been detected in cardiac mitochondria, where genetic and pharmacologic studies show modulation of respiration, mitochondrial Ca^2+^ handling, and reactive oxygen species (ROS) generation, converging on smaller infarcts after I/R or ischemic preconditioning (IPC)(*14, 18, 23, 25, 27, 28, 36, 53-55*). Our work extends this knowledge from isolated cell and injury models to the basal, whole-organ level: by combining comprehensive, non-invasive echocardiography with vascular Doppler and transcriptional profiling, we demonstrate that expression of BK channels is *vital for physiological cardiovascular function* at baseline, and that loss of *Kcnma1* alone is sufficient to produce global cardiac and aortic dysfunction.

### BK channels are required beyond cardioprotection

Prior studies localized adult cardiomyocyte BK predominantly to the inner mitochondrial membrane, defined its molecular composition (including β-subunit interactions), and linked pharmacologic mitoBK activation to I/R protection(*7, 22, 56, 57*), whereas SAN-localized BK channels regulate intrinsic pacemaker rate(*16, 17, 52*). Our data establish that BK channel expression is necessary for physiological left- and right-ventricular performance, diastolic filling (abnormal E/A, E/E′), and aortic outflow, independent of IR injury. Pharmacologic inhibition *in vivo* and in isolated hearts had suggested a role for BK current in heart rate and ventricular function, but the mechanistic locus (SAN *vs*. working myocardium *vs*. vasculature) and the directionality of effects remained unresolved because of drug off-target actions and systems-level interactions. The present genetic evidence in intact animals closes this gap and, together with our chamber-specific and vascular phenotyping, positions BK as regulator of basal cardiovascular function.

Importantly, the above-mentioned findings have often been using the pharmacologic agents. Widely used openers such as NS1619 can modulate non-BK targets (e.g., Ca^2+^-gated Cl^-^ currents and Ca^2+^ handling)(*58-60*), making conclusions drawn from drug studies not fully reliable. Hence our study in a genetic model, studying the function of BK in physiological cardiac function is an important addiction to define the role of BK.

### BK mitochondria crosstalk and its influence on the UCP axis

BK channels in adult cardiomyocytes localize to mitochondria, are targeted by specific C-terminal splice variants, and co-assemble with β-subunits to tune Ca^2+^/voltage gating and mitochondrial Ca^2+^ retention(*7, 18, 19, 25, 26, 54, 55, 61-68*). In keeping with this, our unbiased transcriptomics from *Kcnma1*^*-/-*^ cardiomyocytes revealed differential regulation of pathways central to heart size, cell death, pulmonary hypertension, and arrhythmia. Among mitochondrial genes, all three uncoupling proteins (UCP1/2/3) were downregulated, with validation at the protein level for UCP1 and UCP3. Consistent with established roles of UCPs in attenuating mitochondrial ROS and in tuning substrate oxidation, the hypertrophy-fibrosis-dysfunction upon loss of BK consolidates the predicted roles of BK to bioenergetics and redox control. UCP2/3 induction is an established component of delayed preconditioning in the heart, where they contribute to ROS moderation and improved survival, further supporting their importance to cardiomyocyte stress tolerance.

Mechanistically, our data implicate a retrograde signal from the mitochondrial BK to the to nuclear programming: PGC-1α, a master coactivator of mitochondrial biogenesis and oxidative metabolism (including UCP transcription), is downregulated in BK-null hearts; in parallel, FOXO3a, which cooperates with and transcriptionally induces PGC-1α to orchestrate antioxidant defenses, is dysregulated at the phosphorylation level. It is to be noted that despite UCP suppression, we did not detect broad OXPHOS protein loss, arguing for a selective UCP/antioxidant remodeling rather than a global mitochondrial gene depression. Such selectivity aligns with mitoBK’s known impact on mitochondrial Ca^2+^ and ROS microdomains signals that drive transcriptional programs without necessarily collapsing respiratory-chain abundance.

### β3-adrenergic mediated UCP expression rescues cardiac function in BK mutants

We found that augmenting UCP expression with a β_3_-adrenergic agonist (CL-316,243) normalized ejection fraction in *Kcnma1*^*-/-*^ mice and upregulated UCP1/2 proteins. While the canonical site of β3-AR-induced UCP1 expression is brown/white adipose tissue, this pharmacological method robustly induces UCP1 and thermogenic programs and increases systemic energy expenditure(*44, 45*). This is in line with our observation that boosting UCPs can compensate for upstream BK deficiency to restore cardiac function. B3-ARs are present in the mammalian heart and have complex metabolic and contractile effects(*69*); our data suggests that targeted study of β3-AR-UCP coupling in cardiomyocytes can provide novel therapeutic avenues.

### Limitations and future directions

First, although our microarray and immunoblotting converge on a BK to PGC-1α/FOXO3a to UCP axis, the *proximal* signaling events linking mitoBK activity to nuclear transcription remain undefined. Elucidating whether mitochondrial Ca^2+^ transients, matrix NAD(P)H/ROS signals, or OMA1/DELE1-like stress axes mediate BK-dependent retrograde communication will be essential. Second, cell type-specific *Kcnma1* deletion and rescue (e.g., cardiomyocyte-restricted mitoBK ablation versus vascular smooth muscle deletion) are required to explain the relative contributions of cardiac versus vascular BK to the reported phenotypes. Finally, while β_3_-agonist driven UCP upregulation rescued function in our model, dose response and tissue selectivity studies are needed to delineate direct cardiac effects from systemic metabolic adaptations and to ensure absence of pro-arrhythmic or maladaptive remodeling in the long term.

## Conclusions

In summary, our genetic, physiological, and molecular analyses establish BK channels as non-redundant factors of basal cardiovascular function and identify a mitoBK to PGC-1α/FOXO3a to UCP pathway as a mechanistic bridge to mitochondrial health.

## Author contributions

S.G.R, N. P., N. J. P., K. S., A. T. H., S. G., S. S., S. K. R., D. P., P. K., S. A., A. K., and H. S. performed experiments. F. A., S. A., and A. K., provided resources and key experimental material. S.G.R. and H.S. obtained funding. S.G.R., N. P. and H. S. wrote the first draft.

## Funding and additional information

S.G.R and H.S are supported by AHA TPA (972077 and 965031, respectively). This work is supported by the National Heart, Lung, and Blood Institute, National Institutes of Health grant (grant nos.: HL157453 and HL179189 [to H. S.]). The content is solely the responsibility of the authors and does not necessarily represent the official views of the National Institutes of Health.

