## Supplementary Information for "BK Channels Orchestrate Cardiac Homeostasis Through Mitochondrial Uncoupling Proteins"

**Supplementary figure 1**

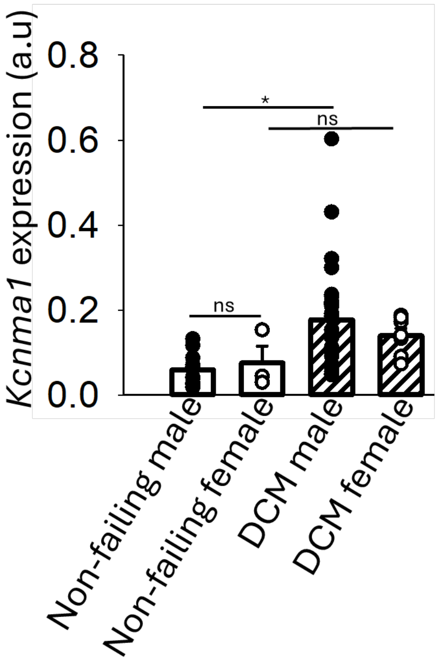


**Supplementary figure 1: BK levels increase in failing human male hearts.** Expression of BK mRNA increases in end stage failing hearts with dilated cardiomyopathy compared to non-failing hearts only in males but not females.

**Supplementary figure 2**

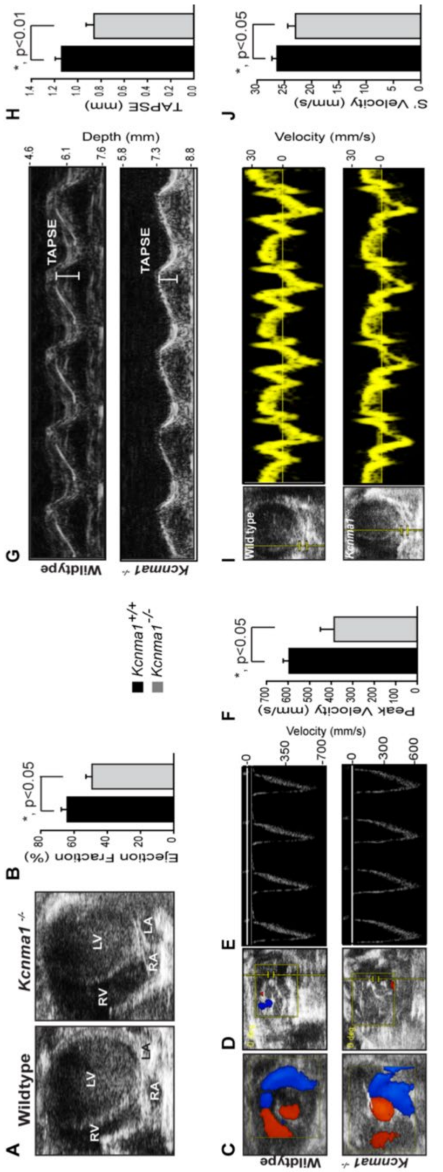


**Supplementary figure 2: . The absence of BK causes right heart dysfunction.**

**A.** Echocardiogram showing RV enlargement in *Kcnma1*$^{-/-}$ compared to WT hearts. **B**. RVEF is also reduced in KO hearts compared to WT hearts. **C** and **D**. Color flow doppler showing blood flow through the Right Ventricular Outflow Tract (RVOT) and through the pulmonic valve. **E** and **F**. Doppler interrogation of peak pulmonary artery flow measured from C and D shows reduced velocity in KO hearts compared to WT hearts. **G** and **H**. TAPSE is also reduced in KO hearts compared to WT hearts. **I** and **J**. Tissue doppler S' is reduced in KO hearts compared to WT hearts.

**Supplementary figure 3**

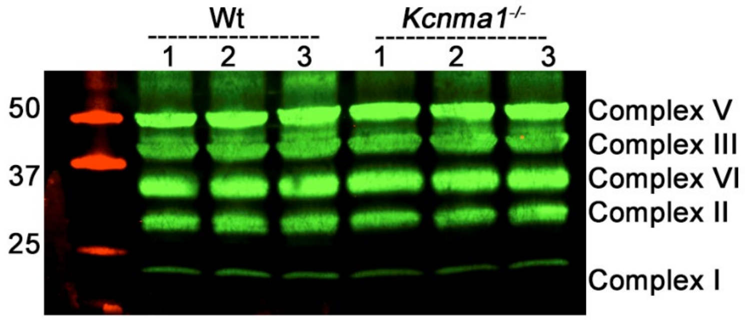


**Supplementary figure 3:** Western blot analysis of mitochondrial complex protein levels in wild type and BK knockout animals. There were no significant differences between the levels of proteins in wild type and BK knockouts.
